# Active exploration generates observer-specific image cues for material identity

**DOI:** 10.64898/2026.11.13.766475

**Authors:** Takuma Morimoto, Ryu Nomachi, Hideki Tamura

## Abstract

When observers interact with an object, each observer generates a different sequence of images. However, which image statistics underlie their perceptual judgments, and whether this self-generated visual input explains what they perceive, remain unclear. We analyzed the first-person recordings of a virtual-reality experiment in which 19 observers judged whether an object was metal or glass while exploring the object with the hand, with the head, with both, and without moving. The object was rendered in two shapes at five levels of physical metallicity and in four illumination environments; every observer saw every stimulus but did so through a different video. Simple image statistics pertaining to the object, to its relation with its surround and to the changes in these properties as the observer explored predicted observers’ responses effectively. Critically, when observers disagreed with regard to the same stimulus, their own videos predicted which observer said “metal” more accurately than chance, and this observer-specific information emerged mainly in the few seconds that preceded the response. The predictive cues identified in this context included the coverage of the optic flow, contrast, highlight coverage, luminance, and the object’s luminance and color relative to its surround; furthermore, their importance varied across observers. Thus, active exploration generates different visual evidence for different observers, and their material judgments follow the evidence that they generate.

## 1. Introduction

Perceiving what an object is composed of is a challenge in image interpretation. The light that reaches the eye depends jointly on the surface, its shape and the illumination, and the visual system must read the material from the image structure that confounds these three aspects (Adelson, 2001; Anderson, 2011; Fleming, 2014, 2017; Anderson & Marlow, 2023; Schmidt et al., 2025). Many image cues have been proposed, such as the luminance histogram, its skewness and its contrast (Motoyoshi et al., 2007; Sharan et al., 2008; Wiebel et al., 2015), the coverage, sharpness and contrast of highlights (Marlow et al., 2012; Marlow & Anderson, 2013), their congruence with shading (Anderson & Kim, 2009; Kim et al., 2011), the cues that indicate metal (Todd & Norman, 2018; Harvey & Smithson, 2021), the specular structure that separates material categories (Schmid et al., 2023), the distortions that a transparent body imposes on its background (Fleming et al., 2011), the specular reflections on its surface (Schlüter & Faul, 2014, 2016), the image of the surroundings, which a refracting body presents inverted and a mirror presents upright (Kim & Marlow, 2016), and the transmitted light and internal reflections that indicate glass (Todd & Norman, 2019, 2020). Because the cues depend on the illumination, perceived gloss is only partly constant across light fields (Fleming et al., 2003; Pont & te Pas, 2006; Olkkonen & Brainard, 2010; Motoyoshi & Matoba, 2012; Chadwick & Kentridge, 2015; Zhang et al., 2019; Morimoto et al., 2023), and perceived roughness (Ho et al., 2006) and the appearance of transparent and translucent materials (Xiao et al., 2014; Gigilashvili et al., 2021) change with the direction of the light. Image-computable models that rely on hand-designed statistics (Nishida, 2019; Cheeseman et al., 2025) or on features that are learned by neural networks (Storrs et al., 2021; Prokott et al., 2021; Prokott & Fleming, 2022; Tamura et al., 2022; Liao et al., 2023; Morimoto et al., 2026) can now predict human material judgments accurately.

In almost all of this work, however, the image has been chosen by the experimenter. Observers view a given picture, and when the stimulus moves, the motion is scripted (Hartung & Kersten, 2002; Doerschner et al., 2011, 2013; Marlow & Anderson, 2016; Kawabe & Kogovšek, 2017; Tamura et al., 2018; Tamura & Nakauchi, 2018) or its range is fixed (Sakano & Ando, 2010; Tani et al., 2013). The same characterization applies in the contexts of motion cues concerning shape (Todd, 2004; Dövencioğlu et al., 2015, 2017) and the mechanical properties of elastic objects, liquids, cloth and breaking objects (Paulun et al., 2017; Schmidt et al., 2017; Schmid & Doerschner, 2018; van Assen et al., 2018, 2020; Bi et al., 2019; Alley et al., 2020; Kaiser et al., 2023). Outside the laboratory, the observer chooses the images. We turn an object in the hand, tilt our head and look again, so that the sequence of images on the basis of which we judge a material is partly of our own making. Such active sensing has been well studied in touch (Gibson, 1962; Lederman & Klatzky, 1987) as well as eye movements and natural behavior (Hayhoe & Ballard, 2005; Najemnik & Geisler, 2005; Yang et al., 2016; Gottlieb & Oudeyer, 2018; Goettker et al., 2025), in which each fixation has been treated as a sample taken to gain information. In material perception, however, the question is more recent. Observers tilt a hand-held display to judge the surface it simulates (Ferwerda & Padhye, 2021), explore differently in situations involving gloss than in those involving lightness (Lin et al., 2025), fixate the diagnostic regions (Toscani et al., 2013, 2019; Metzger et al., 2024) and, in virtual reality, move most when the material in question is ambiguous (Nomachi et al., 2026). Material perception has accordingly been proposed to be an active process (Xiao & Liao, 2025); observers differ in how they weigh color relative to material (Brainard et al., 2018) and may rely on different visual cues (Liao et al., 2022). Materials are a good test case for active exploration: a single image is often ambiguous about what an object is made of, and exploration may be the strategy that observers use to resolve that ambiguity.

The fact that observers explore is thus established. The question that remains unanswered is what they gain from it, including in terms of which visual features they look for and whether the information that they collect directly informs their judgments. Moreover, whether two observers who explore the same object in different ways collect different visual evidence and consequently reach different judgments is unclear. To address this gap, we modeled first-person videos that were recorded in an experiment conducted by Nomachi et al. (2026): the view of every observer was captured throughout a material discrimination task in virtual reality. Nomachi et al. analyzed the responses and the movements of the head and hand, not the videos themselves. Observers judged whether a floating object was metal or glass. The object was rendered at five levels of physical metallicity, in two shapes and four environments, and in separate conditions observers could turn it with the hand, move their head, or both; a fourth condition, in which they could not move, provided a baseline without exploration. Each of the 19 observers judged 120 such trials with exploration (40 trials for each mode of exploration) and 40 without, and for every judgment, we had access to the frames that the observer saw. Crucially, the stimulus was identical for every observer, whereas the video differed. We ask three questions: which image and video features predict material judgments? Can an observer’s own frames explain their response, as tested between observers who saw the same stimulus but disagreed with one another? When does this observer-specific information emerge during exploration?

## 2. Methods

### 2.1 The experiment and its recordings

The data used in this research are first-person recordings that were produced during Experiment 3 from Nomachi et al. (2026). Observers wore a head-mounted display (Varjo Aero, 2880 × 2720 pixels per eye, 115° field of view, 90 Hz) and held a controller in each hand. The scene, which was built in Unity (2022.3.22f1) with the high-definition render pipeline (HDRP), contained a high-dynamic-range illumination map and one target object, which was 6 cm across and floated 50 cm in front of the observer (6.9° of visual angle). The object featured one of two shapes, a relatively smooth shape ("smooth") or a more irregular shape ("bumpy"), both of which were generated by fractal modulation of the surface of a sphere (Tamura et al., 2018). It was rendered with the HDRP Lit shader (smoothness 1.0, index of refraction 1.424, thickness 0.8) with the shader’s metallic parameter was set to 0, 0.25, 0.5, 0.75 or 1 (Figure 1a). We refer to this parameter as the physical metallicity of the object. It sets the balance between the refraction of the background, which dominates at 0, thus making the object a glass-like body, and the reflection of the environment, which dominates at 1, thus making the object polished metal; the intermediate levels are physically ambiguous mixtures. The four environments were an outdoor scene featuring the sun ("Sky"), an indoor room ("Studio"), a black-and-white checkerboard ("Check") and a low-contrast checkerboard of two gray shades ("Gray"; Figure 1b,c).

**Figure 1.**
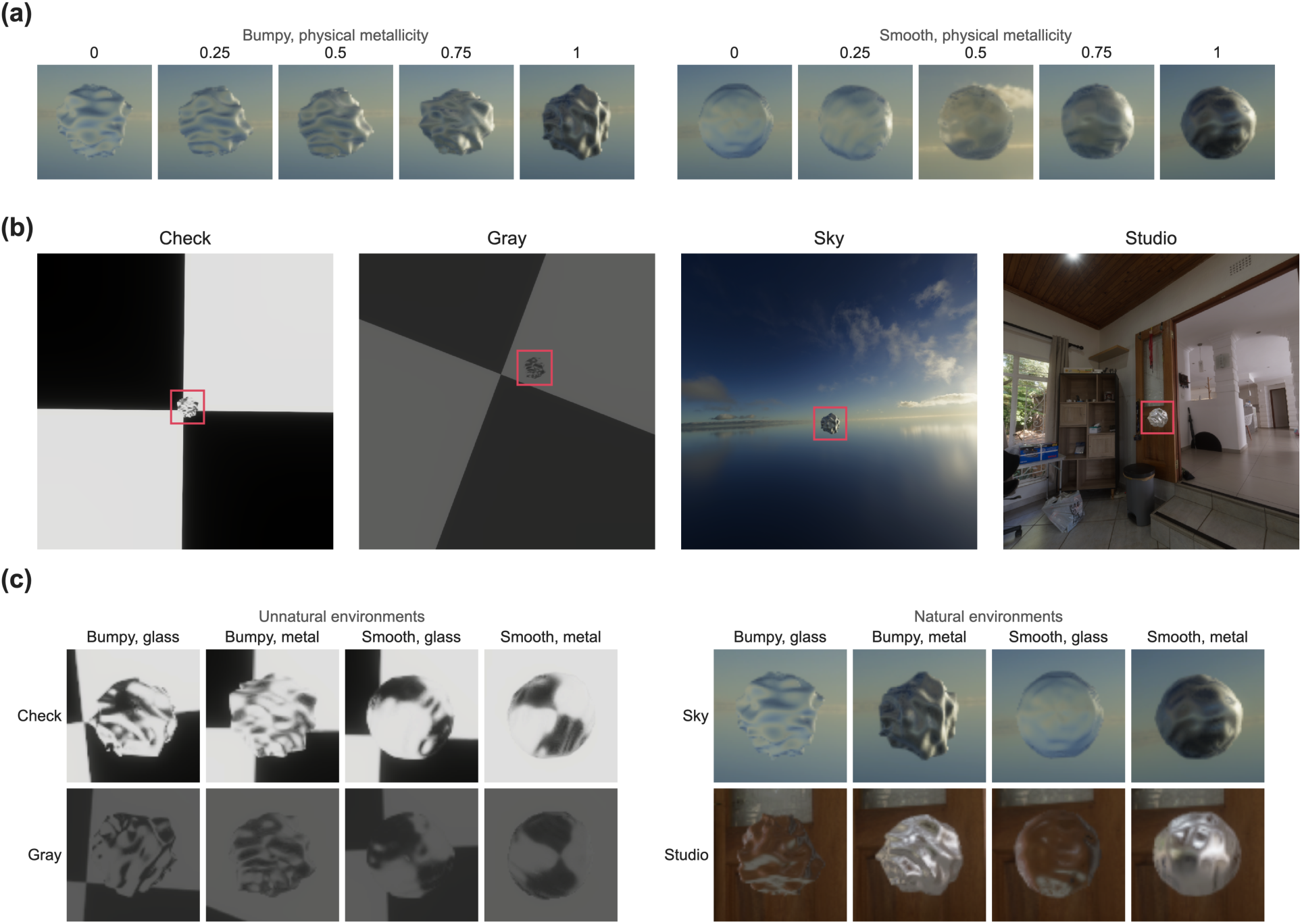
Stimuli, recordings and model inputs. (a) The object at the five levels of physical metallicity for the two shapes under the Sky illumination, illustrated as the analyzed patch. (b) First-person view of one observer in each illumination environment (bumpy object, physical metallicity 1). The box indicates the patch analyzed: the object and a margin of a quarter of its size on each side. (c) The patch for the two shapes at the two ends of physical metallicity in the four environments.

In each trial, the observer viewed the object, explored it freely in the three exploration conditions with no time limit, and pressed a button to report whether it was metal or glass. The experiment included four conditions that involved 40 trials each, namely, one condition per combination of head tracking and controller tracking: (i) with the head only, in which the viewpoint followed the head; (ii) with the hand only, in which the observer could grasp, move and rotate the object; (iii) with both, in which both modes were available; and (iv) in the static condition, in which nothing moved. Within a condition, the 40 trials (i.e., 5 levels of physical metallicity × 2 shapes × 4 environments) were presented in random order, and the order of the conditions was counterbalanced. Throughout each trial, the software captured the image that was shown to the observer, which was cropped to a 720 × 720 pixel square around the center of the view, approximately every 17 ms (median 16.7 ms). Each trial thus yielded a video extending from the moment the object appeared to the response, accounting for 33 to 2,490 frames (median 258) or 0.5 to 41 s of exploration (median 4.4 s).

### 2.2 Observers

Twenty students at the Toyohashi University of Technology, who had normal or corrected-to-normal vision and who were naive to the purpose of the study, participated; one individual was excluded from the study by Nomachi et al. (2026) because the images could not be brought into focus for that observer, thus leaving a total of 19 participants. The study was approved by the Committee for Human Research of the Toyohashi University of Technology, and all observers provided written informed consent.

### 2.3 The videos: Objects, surrounds and sampling

From each frame, we cut two regions. The object crop is a square window of fixed size, which is identical for every frame (the size was sufficiently large to contain the object in 99% of all frames) and centered on the object but resized to 224 × 224 pixels. The object itself is always 6 cm in size, but observers could bring it closer with the hand or the head. Because the window does not scale, a closer object appears to be larger in the crop; thus, the object’s area in the crop records its distance. The patch, from which the statistics used to compare the object with its surround are computed (section 2.4), follows a different rule: it scales with the object, extending a quarter of the object’s size beyond its bounding box on each side (see the box in Figure 1b and the patches in Figure 1a,c), such that the object always occupies approximately the same fraction of the patch and a fixed amount of surround is included. We included the surround because a glass object presents a refracted image of the background behind it (Fleming et al., 2011; Kim & Marlow, 2016), whereas a metal object reflects the environment around it (Fleming et al., 2004; Tamura et al., 2018, 2022); how the object is related to its surround therefore serves as a cue in its own right, alongside the cues within the object. Frames in which the object had left the view were skipped (accounting for 352 of 296,662 sampled frames; in 45 of 2,280 trials); in every other frame, the object covered at least approximately 3,000 pixels of the crop, which was sufficient for every statistic.

### 2.4 Image and video statistics

We used three families of statistics. Two of these families, i.e., “object" and “relational", describe each frame and are summarized across frames; the third family, i.e., “motion”, describes the change between frames. Each statistic is defined in Figure A1 and Table A1 in the Appendix.

In plain terms, the object statistics characterize the intuitive properties of the object: how bright it is, the extent to which its brightness is spread over its surface, how much of it is covered by highlights, how colorful it is, how large it appears to be and how much it changes from one frame to the next. The relational statistics focus on how bright and colorful the surround is and compare the object with it. The object statistics are simple image-computable measures that are motivated by previous work on lightness, gloss, metal and transparency (Motoyoshi et al., 2007; Sharan et al., 2008; Marlow et al., 2012; Wiebel et al., 2015; Todd & Norman, 2018, 2019; Harvey & Smithson, 2021) and are computed over the object’s pixels; the relational statistics are computed over the patch. Each frame includes 15 object quantities: the level and spread of luminance and the shape of its distribution (skewness and kurtosis), its contrast, the coverage and contrast of highlights, the amount of fine edge structure (edge energy), the fraction of dark pixels, colorfulness (chroma), the object’s area in the crop and the frame-to-frame change. A total of 9 relational quantities are included: the mean, spread and maximum of the surround’s luminance, its contrast and its chroma, and the ratios of the object to its surround in mean luminance, contrast, maximum luminance and chroma. Each quantity is a time series over the trial, which is summarized by reference to its mean, standard deviation, and maximum, as well as the difference between its means over the second and the first half of the trial (and, for four quantities, by how similar consecutive values are, the lag-one autocorrelation). The 15 object quantities thus yield 15 × 4 summaries + 3 autocorrelations = 63 object statistics, and the 9 relational quantities yield 9 × 4 summaries + 1 autocorrelation = 37 relational statistics: 100 image statistics in total. None of these quantities is specific to metal or glass by design; the summaries over time allow a model to use how the image changed as the observer moved.

A third family is adapted from the optic-flow cues pertaining to shininess identified by Doerschner et al. (2011), which focus on how image points move from one frame to the next. We placed a grid of points on the object every 6 pixels and tracked each point to the frame two captures later, a separation of approximately 34 ms (pyramidal Lucas–Kanade tracking; Lucas & Kanade, 1981; Bouguet, 2001), keeping a point only if tracking it back led to a return to within 1 pixel of its start. For each frame pair, we computed the coverage (the fraction of points that could be tracked), its change when the distance between the frames is doubled (both adapted from Doerschner et al., 2011), the divergence of the dense flow field (Farnebäck, 2003; this aspect indicates whether the motion spreads apart or converges, as a fraction of the object and as a mean), the 3-D shape reliability (i.e., the fraction of tracked points whose motion is consistent with a rigid object turning, as inliers of a fundamental matrix; Fischler & Bolles, 1981) and the mean speed. These quantities were summarized over the trial as described above. The divergence fraction and 3-D shape reliability are undefined in 1 to 2% of the videos; these cases featured an insufficient number of trackable points and were left out, thus leaving 4 quantities × 4 summaries = 16 motion statistics.

In total, 116 statistics (63 object, 37 relational and 16 motion) were used. The object and relational image statistics describe each frame and are summarized over the trial. We analyzed up to 150 frames per trial, which were evenly spaced over the recording (a total of 31% of the videos had 150 frames or fewer and were used whole); the alternative use of 300 frames yielded practically the same values (correlation *r* > 0.99 for 90% of the statistics). The motion cues, for which the time between frames itself was meaningful, used a fixed interval of two captures instead. In the static condition, the image did not change within a trial (it was rendered from the point at which the observer’s head was located when the trial began), so each trial consisted a single frame, which was described solely by the per-frame quantities indicated above, accounting for 23 in total, without the summaries over time and the frame-to-frame change.

### 2.5 Models and their evaluation

Each video was represented by 116 object, relational, and motion statistics. We used logistic regression to predict the observer’s binary response ("metal" or "glass") on the basis of these statistics. The weighted sum of the statistics before conversion to a response probability was taken as the predicted metallicity of the video (i.e., the log-odds of a “metal” response), in which context higher values indicated a stronger prediction of “metal”. Each statistic was standardized within the observer before fitting, such that the model used trial-to-trial variation rather than differences in the overall level of a statistic between observers. We used L2 regularization (C = 0.1, selected by grid search).

Models were evaluated using leave-one-observer-out cross-validation: each model was fitted to the videos and responses of 18 observers and used to predict the responses of the remaining observer; this approach was repeated for all 19 observers. For the disagreement analysis presented in section 3.4, in which videos obtained from different observers who viewed the same stimulus were compared, we instead used leave-one-object-out cross-validation. For each of the 40 objects (shape × physical metallicity × environment), the model was fitted on the basis of videos and responses for all observers for the other 39 objects across all conditions. Thus, no response to the object being tested was used for fitting, and all observers’ videos for that object were evaluated by the same model.

Prediction accuracy was quantified by the area under the receiver operating characteristic curve (AUC). Unless otherwise stated, the AUC was calculated by reference to observers’ responses, not the physical metallicity of the stimulus. This factor therefore measures how well predicted metallicity distinguishes trials that were judged as “metal” from those that were judged as “glass”. Equivalently, the AUC is the probability that, given one trial that was judged as “metal” and one that was judged as “glass”, the model assigns a higher predicted metallicity to the former. An AUC of 1 indicates perfect prediction and a 0.5 chance. The procedure leading from an example video to its predicted metallicity and AUC is illustrated in Figure 2a–c.

**Figure 2.**
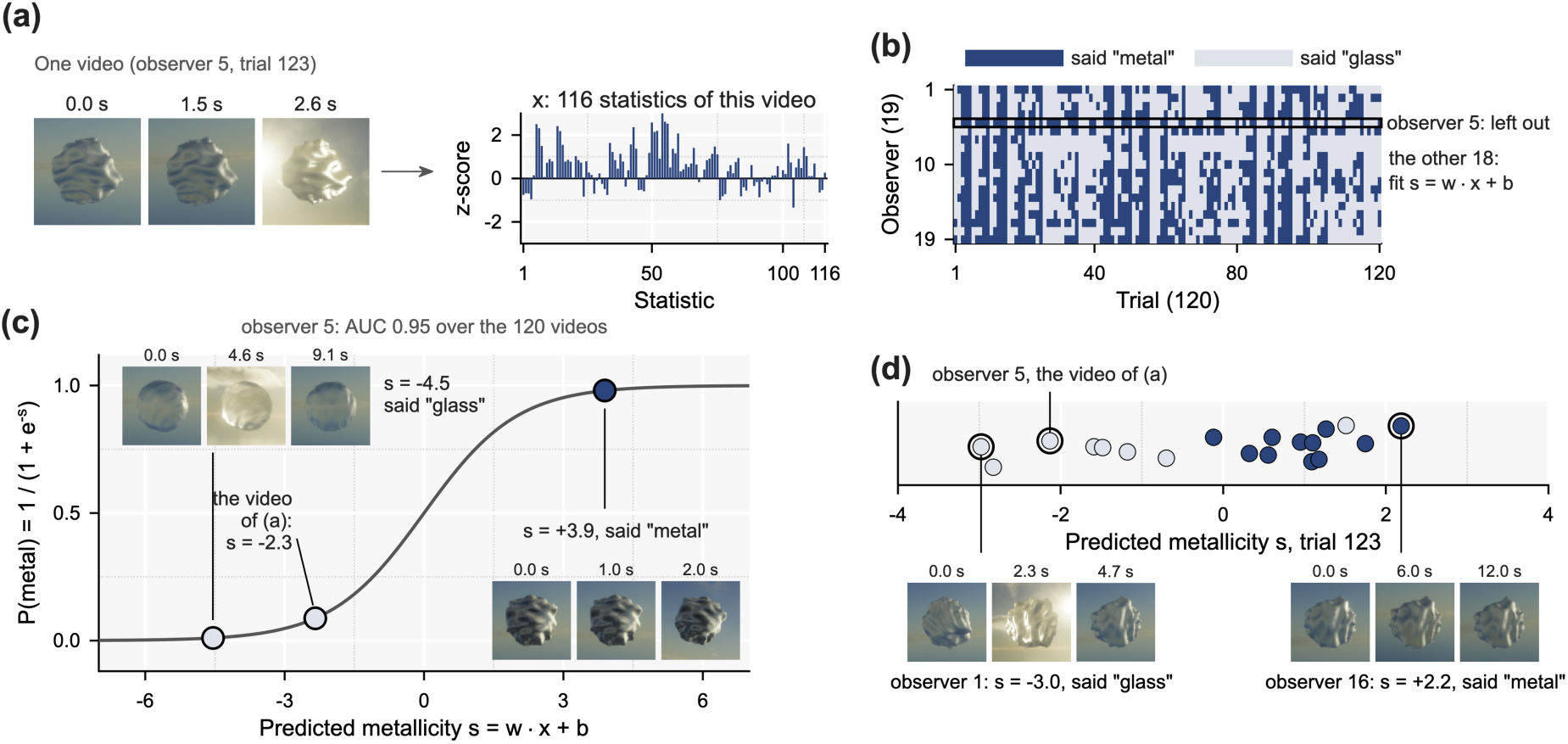
How the observer’s-own-video model is developed and tested. (a) One video (observer 5, trial 123: bumpy object, physical metallicity 0.5, Sky, head and hand) becomes a list x of 116 statistics, each of which is standardized within the observer. (b) Responses of the 19 observers to the 120 trials. To predict observer 5, their row is left out, and a logistic regression s = w·x + b is fitted to the 2,160 videos and responses of the other 18 observers. (c) The fitted model scores observer 5’s 120 videos, which it has never seen; s is the predicted metallicity, which is the log odds of a “metal” response. (d) Individual variation in predicted metallicity: the videos of the same trial (trial 123) for 19 observers, scored by a model fitted to the videos of the other 39 objects (the model of the disagreement test, section 3.4), including the lowest- and the highest-scoring video; the color indicates each observer’s response.

We used two reference predictors to interpret the model AUC. Physical metallicity provides the performance expected from a predictor that knows the metallicity parameter but not the observer. Agreement among observers serves as a second reference: for each trial, we used the proportion of the other 18 observers who responded “metal” to predict the held-out observer’s response. Because this proportion was estimated from a finite sample, it provides a lower estimate of the agreement ceiling; including the held-out observer’s own response provides an upper estimate (cf. the noise ceiling of Nili et al., 2014). We refer to the range between these estimates as the ceiling.

## 3. Results

### 3.1 Behavior

Nomachi et al. (2026) analyzed the behavior observed in this context in full; we provide only a brief overview here. All tests presented in this paper are two-tailed, and the 19 observers were used as the unit. Effect sizes are Cohen’s *d* for paired and one-sample comparisons (the mean difference divided by the standard deviation of the differences) and Kendall’s *W* for Friedman tests. The proportion of metal responses increased with physical metallicity, as expected (Figure 3a, b, c). The environment mattered (Figure 3a): under the natural maps, objects at physical metallicity 1 were nearly always called metal (0.98 Sky, 0.91 Studio), whereas under the artificial maps, the curve was shallow. The bumpy shape was called metal more often at every level (Figure 3b). Exploration entailed that the responses followed physical metallicity more closely (Figure 3c). Without exploration, a glass object (physical metallicity 0) was called metal in 0.23 of the trials against 0.09 with exploration (paired *t*(18) = 3.8, *p* = 0.001, *d* = 0.87). Observers looked at the fixed image for 3.0 s on average and explored for 5.6 s (Figure 3d; paired *t*(18) = −5.5, *p* < 0.001, *d* = −1.27), doing so for the longest when the hand was available (6.4 s hand only, 5.7 s head and hand, 4.6 s head only; Friedman χ²(2) = 10.2, *p* = 0.006, Kendall’s *W* = 0.27). When the hand was available, observers also looked longer when the material was ambiguous: the duration peaked at the intermediate levels of physical metallicity, whereas with the head only or without exploration, it did not vary substantially with the level.

**Figure 3.**
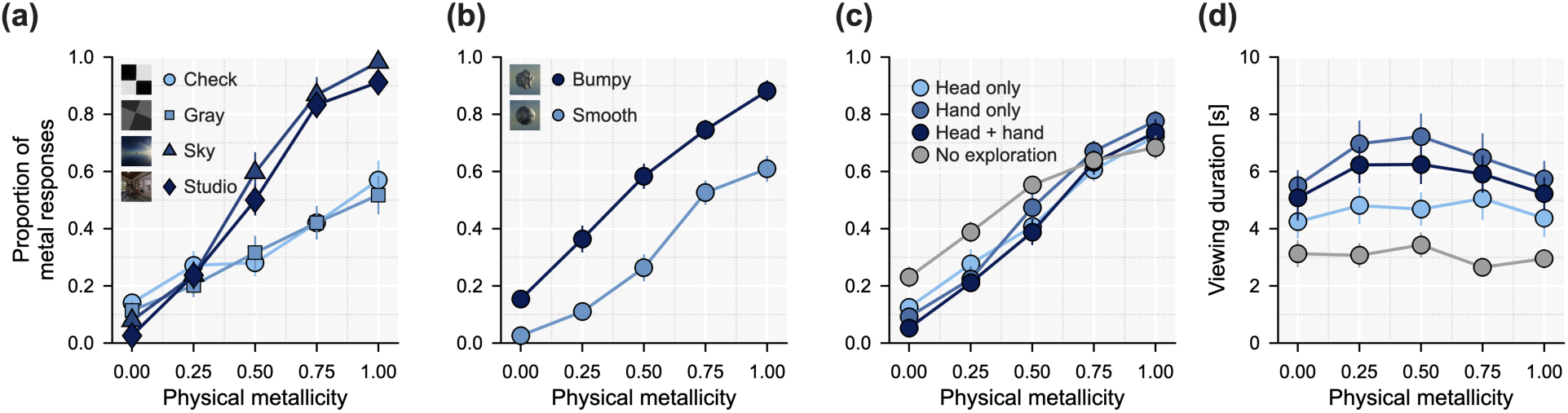
The observers’ responses. (a) Proportion of metal responses against physical metallicity for the four environments and (b) for the two shapes over the three exploration conditions; (c) for each condition, including the condition without exploration. (d) Viewing duration for each condition. Symbols are means over the 19 observers of each observer’s mean; error bars are standard errors across observers.

### 3.2 Predicting the responses from the video

Figure 4a compares three predictors on held-out observers, to whom the model was never fitted, over all trials with exploration. The 116 statistics extracted from the observer’s own video (the first box) reached an AUC of 0.880 (SD 0.063 across observers), up to the ceiling that the observers themselves set, as indicated by the gray band in Figure 4 (section 2.5): its upper bound (0.892) is the most that a model that read only the stimulus could reach, and the statistics achieved this bound (*t*(18) = −1.51, *p* = 0.15, *d* = −0.35). Whatever the observers shared, the statistics captured. Much of this prediction relies on a small number of combinations of statistics. When the 116 statistics were compressed into principal components, combinations of statistics that capture most of their variation (computed on the 18 fitted observers) yielded an AUC of 0.59 with 2 components, 0.74 with 5, 0.83 with 10, 0.86 with 20 (which retain 86% of the variation; the second box of Figure 4a) and 0.87 with 30, against 0.88 with all 116 (Figure 4b); twenty components already convey most of the prediction. Physical metallicity alone (the third box) yielded a value of 0.790 (SD 0.059), which was well below the statistics (paired *t*(18) = 5.08, *p* < 0.001, *d* = 1.17).

**Figure 4.**
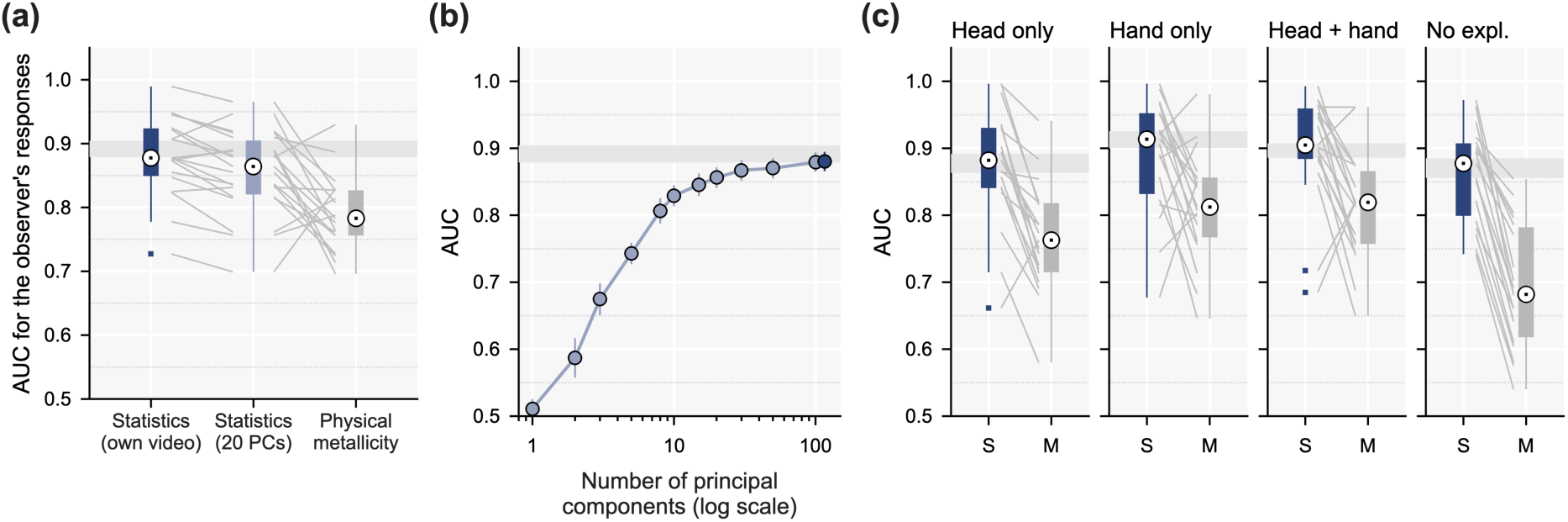
Predictors of the observers’ responses. (a) For each observer, the AUC of three predictors for that observer’s 120 responses: the image statistics of the observer’s own video; the same statistics reduced to their first 20 principal components; and physical metallicity. The video models are fitted to the other 18 observers (leave-one-observer-out). The gray lines connect the values associated with the same observer. The gray band is not a model but rather the ceiling. (b) The AUC of the own-video model against the number of principal components to which the statistics are reduced on a logarithmic axis (mean ± standard error across observers); the rightmost point is the model that includes all 116 statistics. (c) As (a) for the trials in each exploration condition (i.e., head only, hand only, head and hand) and, rightmost, the condition without exploration (S, statistics; M, physical metallicity).

The result is broken down by condition, as illustrated in Figure 4c. With the head only, the statistics gave an AUC of 0.874; with the hand only, the corresponding value was 0.890; and with the head and hand, it was 0.902, thereby reaching the upper bound of the ceiling (0.897; *t*(18) = 0.38, *p* = 0.71, *d* = 0.09). Without exploration (rightmost), the 23 statistics of the single image each observer saw yielded a value of 0.862, slightly below the ceiling for that condition (0.876 to 0.898), and a model fitted only to the moving videos (their per-frame means), when applied to the static image, yielded a value of 0.849. The conditions exhibited few differences (Friedman χ²(3) = 4.7, *p* = 0.19; Kendall’s *W* = 0.08; no pairwise difference reached the level of significance), but they appeared in a consistent order: the statistics’ predictive ability was best when the hand was available and lower when the head only was used or in the condition without exploration. The hand offers the observer the most freedom to choose among different views, and the order suggests that the freer the exploration is, the more the video that an observer generated conveyed the evidence that observer’s responses followed: exploration would then be the acquisition of evidence specific to the observer, and the freer the strategy is, the more informative the views that it acquires are.

The motion statistics contributed to this prediction: in their absence, the 100 image statistics yielded a value of 0.863 (Table A2).

### 3.3 Which statistics convey the information

The previous section revealed that the 116 statistics (or a subset thereof) collectively predicted observers’ responses well. We next considered how informative each statistic was on its own. For each statistic, we computed the AUC against each observer’s 120 responses and then averaged the AUC across observers. Each of the 116 statistics, that is, each summary across frames (mean, *SD*, max, or second half minus first half) of each per-frame quantity, was tested separately; accordingly, several summaries of one quantity could appear among the best. The eight statistics that best predicted “metal” (highest mean AUC) and the eight statistics that best predicted “glass” (lowest mean AUC) are illustrated in Figure 5. Metal responses were associated with high coverage of the optic flow, the fraction of points on the object whose motion could be tracked from one frame to the next (AUC 0.73 for its mean, 0.71 for its maximum), a maximum luminance that exceeded that of the surround (0.68), high object RMS contrast (0.68), high maximum luminance (0.67), high edge energy (0.66), and positive luminance skewness (0.64). In contrast, glass responses were associated with an object that was approximately as colorful as its surround (AUC 0.30 for the ratio of object to surround chroma), greater highlight coverage (0.32), and, more weakly, greater variation in the luminance distribution over the course of the trial (0.40 for the SD of skewness). Thus, trackable image motion and sparse, bright, high-contrast image structure tended to signal metal, whereas objects that were filled with light and took on the color of their surroundings tended to signal glass; this characterization is largely consistent with known cues concerning gloss and transparency (Motoyoshi et al., 2007; Marlow et al., 2012; Tamura et al., 2018, 2022).

**Figure 5.**
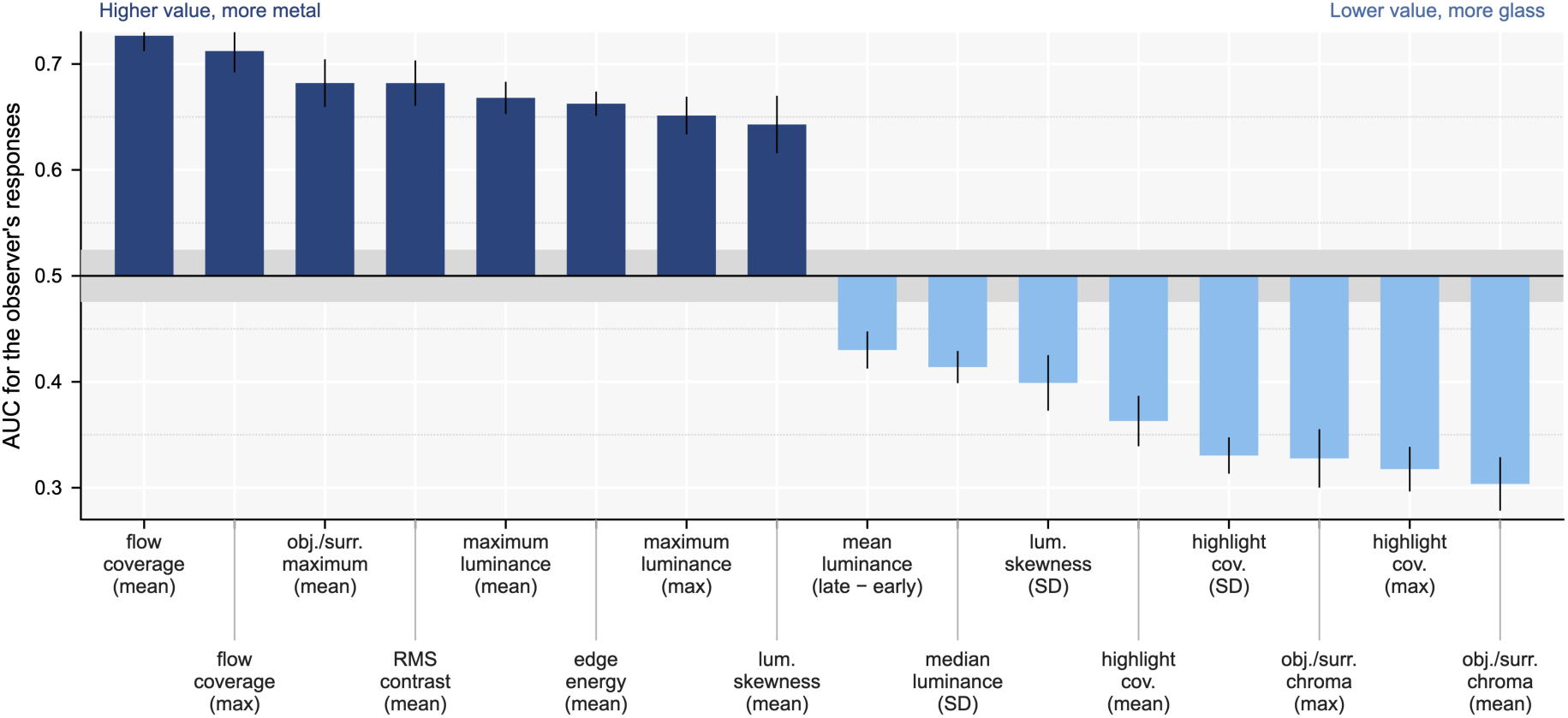
The statistics that predict the responses. The eight statistics for which higher values were most closely associated with “metal” (left, dark) and the eight for which lower values were most closely associated with “glass” (right, light), ranked by their mean AUC. The bars are the AUCs of the statistics (standardized within the observer) against the observers’ responses, including the mean over the 19 observers, alongside the standard error across observers. Values above 0.5 indicate that a higher value was associated with a metal response, whereas values below 0.5 indicate that a higher value was associated with a glass response; the gray band is the range (± 2 *SD*), which is obtained by chance when each observer’s responses are shuffled across their trials (300 shuffles). Each of the 116 statistics, accounting for a summary over the trial (mean, *SD*, max, or second half minus first half) of a per-frame quantity, was tested separately.

Did observers rely on the same statistics? For each observer, the AUC of every statistic across their 120 trials is illustrated in Figure 6; the best predictor of “metal” is presented above each column and the best predictor of “glass” is presented below it. The directions were largely shared: for a typical observer, nearly all of the 16 statistics in Figure 5 (magenta in Figure 6) pointed in the same direction as was the case for the group (median 15 of 16; minimum 10).

**Figure 6.**
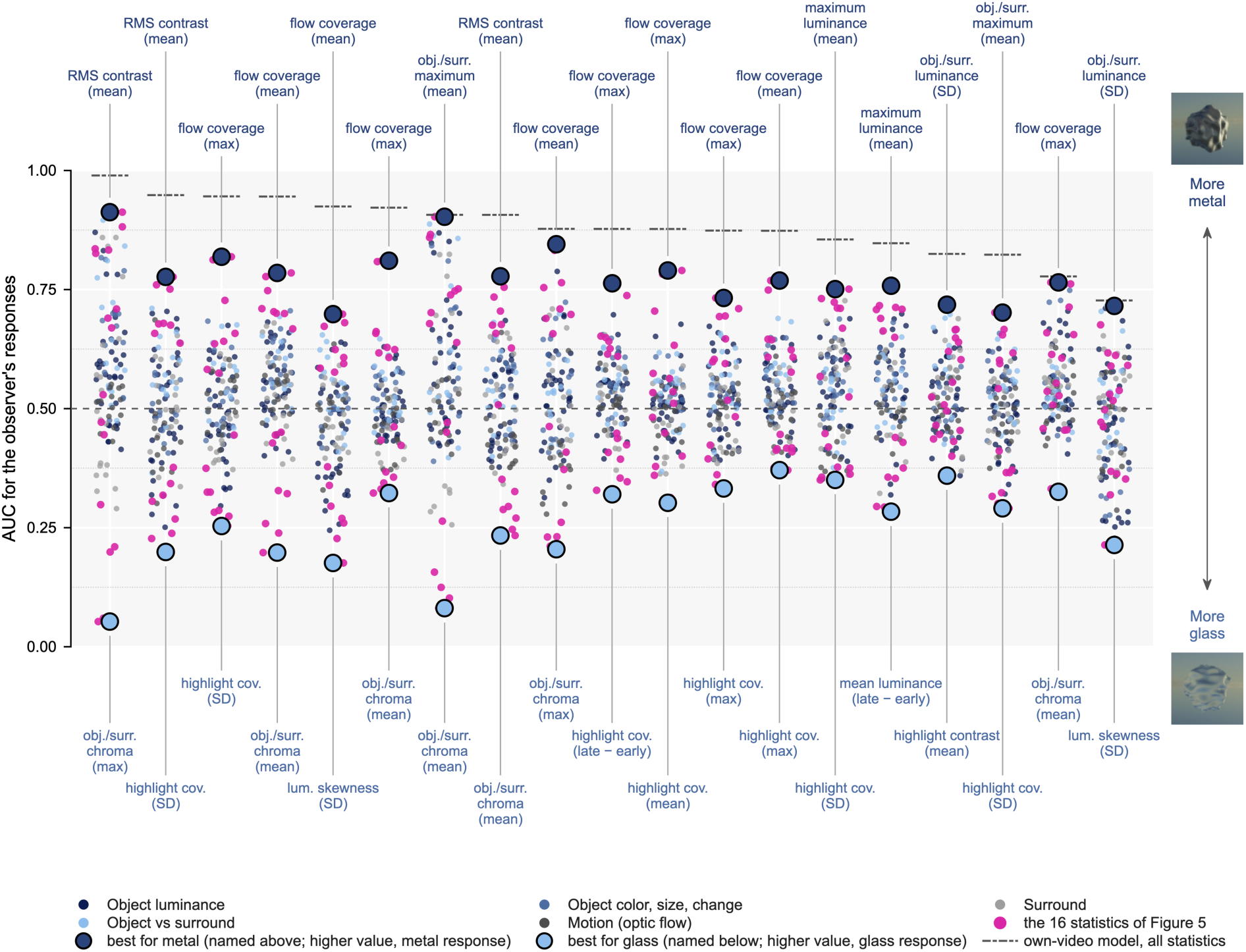
The statistics that predict each observer’s responses. For each observer (columns), the AUC of every statistic over that observer’s 120 trials was as follows: values above 0.5 indicate that a higher value of the statistic corresponded to a metal response, whereas values below 0.5 indicate that a higher value of the statistic corresponded to a glass response. Dots are colored according to the family of the statistic, in which context the 16 statistics presented in Figure 5 are in magenta; the large dark symbol is the statistic that best predicted “metal”, and the large light symbol is the statistic that best predicted “glass”. The dash-dotted line is the leave-one-observer-out AUC of the own-video model, which uses all the statistics.

The best predictors, however, differed across observers: 6 different statistics were best for “metal” (most often the maximum of the flow coverage, accounting for 6 observers) and 9 different statistics were best for “glass” (most often the mean object-to-surround chroma ratio, accounting for 5 observers). The best statistic reached AUCs of 0.70–0.91 for “metal” and 0.63–0.95 for “glass” (expressed as 1 − AUC), thereby approaching the model that used all statistics for some observers (dash-dotted lines). However, the best statistic was rarely much better than the other statistics: for a typical observer, 4 statistics were within 0.05 of the AUC. Thus, observers generally had several informative cues available that pointed in similar directions but differed in how strongly they relied on each cue.

### 3.4 The disagreement test: The observer’s own video explains the observer’s own response

The preceding sections demonstrate that the statistics for the observer’s own video predict the responses as a whole. Some of what these statistics predict, however, is shared by all observers. A diagnostic test would involve asking whether the own video adds something that is specific to the observer. Here, we focused on trials in which two observers explored the same object differently and reached different decisions.

For each trial, we took every pair of observers in which one said “metal” and the other responded “glass” (6,022 pairs over the 120 trials) and asked whether the model attributed the higher predicted metallicity to the video from the observer who said “metal”. For this test, the model is fitted without the trial (Figure 2d), such that it has seen no response to the trial it scores, and all videos of a trial are scored by reference to the same model.

An example disagreement trial is illustrated in Figure 7a: observers 14 and 10 explored the same smooth object but provided different responses. Observer 14 moved the head to look down on the object from above, which situated the object against the dark lower part of the environment (4.1–4.9 s in Figure 7a); from that perspective, the object appeared dark with a few bright reflections, and the observer said “metal”. Observer 10 instead brought the object in front of the bright sky near the sun (1.4–1.8 s), where the view was filled with light, and responded “glass”. Across all 6,022 such pairs, the statistics of each observer’s own video identified the observer who said “metal” with an AUC of 0.667 (Figure 7b, leftmost), i.e., far above the shuffled baseline (0.500 ± 0.017 over 300 shuffles; *z* = 10.0, *p* < 0.004). Shuffling led each response to be reassigned to another observer’s video involving the same stimulus, thus breaking the link between what an observer saw and what they reported while ensuring that the stimulus remained fixed. The near-chance shuffled AUC therefore demonstrates that the predictive information was specific to the video that was generated by each observer. For each observer, the AUC was 0.667 ± 0.011 (mean ± standard error), which was greater than 0.5 for all 19 observers (*t*(18) = 15.26, *p* < 0.001, *d* = 3.50). The effect was general (Figure 7b, others): above chance in every environment, from 0.626 under Gray to 0.705 under Check (*z* ≥ 4.3), and for both shapes (0.669 bumpy, 0.663 smooth).

**Figure 7.**
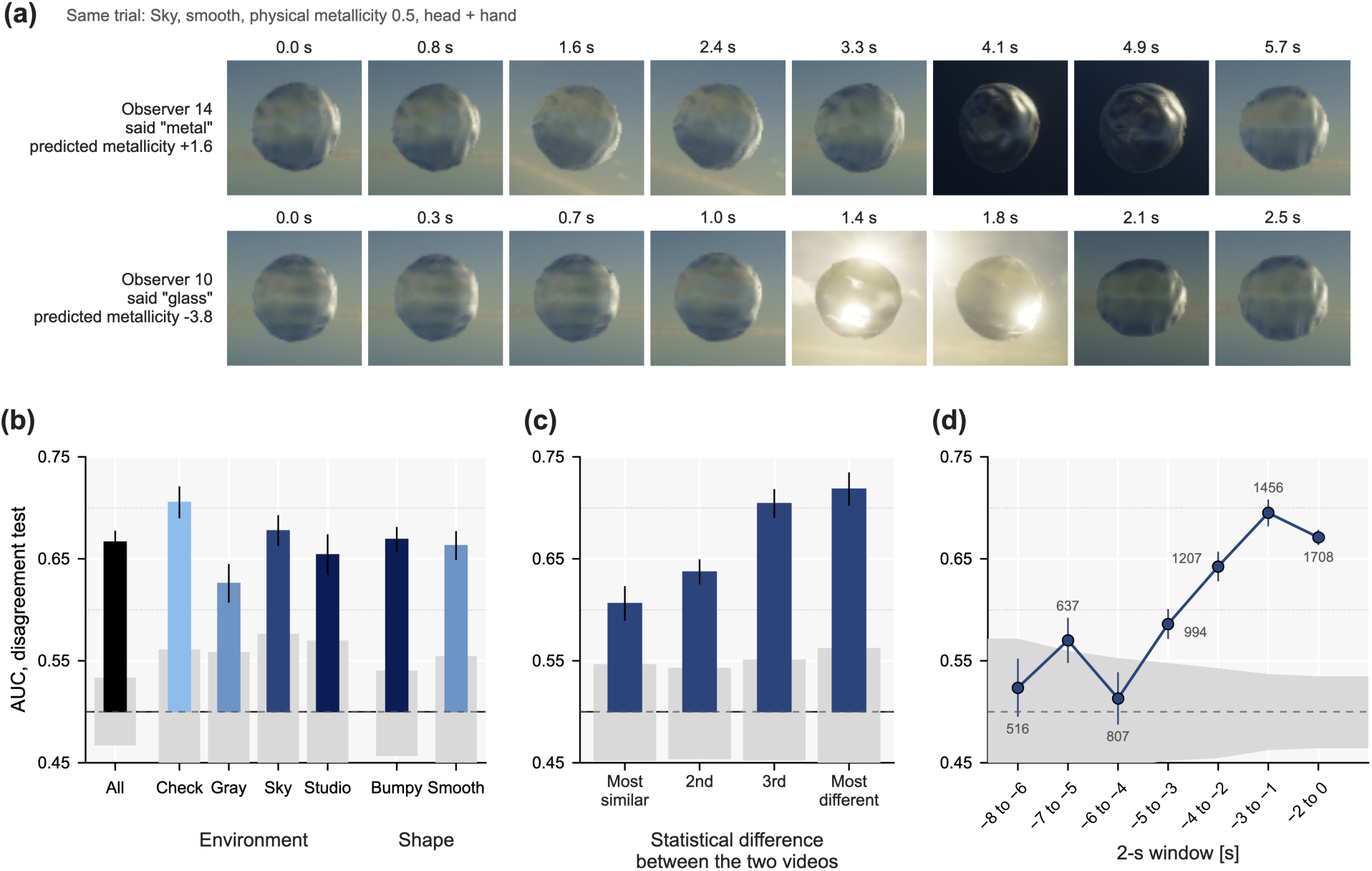
The observer’s own video explains the response beyond the stimulus. (a) Two observers who viewed the same trial but disagreed: eight frames from each video and the predicted metallicity that the model attributed to each. (b) The AUC of the disagreement test over all disagreeing pairs (left) and for the pairs of each environment and of each shape. (c) For the pairs that were split into quarters according to how different the two videos were, the distance between them in the space of the statistics. (d) When the model read the statistics of a 2-s window of each video, it moved back from the response in steps of 1 s; the tick labels provide each window in seconds relative to the response, and the number beside each point is the number of videos that cover the window. In (b) and (c), bars start at 0.5, chance; gray bands are ± 2 SD of the permutation null of each group or window (the responses reassigned at random among the observers of each trial, 300 times); error bars are standard errors across observers of the per-observer AUC (in (d), of the observers with at least 20 disagreeing pairs in the window).

An AUC of 0.67 in the disagreement test is reliable but modest, and we have reason to expect it to be. The test counts every pair of observers who disagreed, including the many pairs whose two videos were nearly identical; such pairs cannot be distinguished by any model and pull the AUC toward 0.5. Therefore, as illustrated in Figure 7c, the pairs were split into quarters in accordance with how different the two videos were. The difference is the Euclidean distance between the two videos’ scores on the 40 principal components that account for 95% of the variation of the 116 statistics (standardized within observers). In the quarter of videos that were most similar, the own videos predicted little as expected (0.606, *z* = 4.5). The AUC then increased through the middle quarters (0.637 and 0.704) to 0.718 in the quarter of videos that were most different (*z* = 7.1). Thus, the more different the evidence that the two observers generated, the more often their own responses followed it.

One might worry that the advantage of the observer’s own video reflects how long the observer explored rather than what the observer saw: the statistics of a longer video may differ for that reason alone. However, duration does not explain this advantage. Alone, this factor exhibited only weak predictive ability (AUC 0.563 in the disagreement test; not shown in Figure 7).

To determine when this information emerged, we computed the same statistics within a 2-s window and moved the window backward from the response in 1-s steps, namely, from the final 2 s to 8–6 s before the response, repeating the disagreement test for each window (Figure 7d). Each window thus contained the same duration of video, and only videos that were sufficiently long to cover it were included (1,708 videos for the final window and 516 for the earliest window). Far from the response, the windows conveyed little information: between 8 and 4 s before the response, the AUC remained near chance (0.524, 0.570, and 0.513; *z* = 0.7, 2.3, and 0.5, respectively). It subsequently increased to 0.586 at 5–3 s and 0.642 at 4–2 s, reaching a peak of 0.695 at 3–1 s before the response (*z* = 10.5; in comparison with the preceding window across observers: paired *t*(18) = 5.54, *p* < 0.001, *d* = 1.27). In the final 2 s, the AUC decreased slightly to 0.671 (in comparison with the peak: *t*(18) = 2.42, *p* = 0.026, *d* = 0.56). This decrease is of interest: observers may have made up their minds by that time, such that subsequent exploration would convey less additional information. Thus, the information that distinguishes observers emerges mainly in the few seconds that lead up to their response, peaking shortly before the response itself, as illustrated in Figure 7a.

## 4. Discussion

We set out to model which image features observers use to distinguish metal from glass when they view an object freely in virtual reality. One hundred and sixteen simple statistics of the images that each observer saw and of the motion between them, which describe the luminance, color and structure of the object, the object in relation to its surround and the optic flow, were sufficient. These statistics predicted an observer’s responses well in every condition, and twenty of their principal components performed nearly as well. The statistics that carried the prediction were the known cues concerning gloss and transparency, and the observers read the same kinds of evidence but weighted them differently.

Our second question was whether active exploration shapes these judgments through the images that the observer generates. The results indicate that it does. In the trials in which the two observers disagreed regarding the same object, the statistics of each observer’s own video predicted which of the two observers responded “metal”. Because both observers saw the same object, the difference pertained to the views that each observer had generated. This observer-specific information was not spread evenly throughout the trial; it was near chance until approximately 4 s before the response and peaked between 3 and 1 s before the response.

The statistics that carried the information are largely consistent with known cues concerning metal and glass. As a mirror-like object turns, reflections move across its surface, and highlights appear, grow and disappear (Doerschner et al., 2011; Dövencioğlu et al., 2017; Tamura et al., 2018), whereas a transparent body transmits light and can reveal itself through distortions of the background (Fleming et al., 2011; Kawabe & Kogovšek, 2017; Todd & Norman, 2019) and through the contours that are produced by the reflections inside it (Todd & Norman, 2020). Thus, an observer who turns the object toward the sun, as illustrated in Figure 2a, produces views in which it is filled with light, increasing highlight coverage and reducing contrast, both characteristic of a refracting body; in turn, views against a darker surround produce sparse highlights and the positively skewed (Motoyoshi et al., 2007) and high-contrast (Marlow et al., 2012) structure that is associated with glossy surfaces. The strongest single cue for metal was the coverage of the optic flow, the fraction of the object’s image that could be tracked from one frame to the next (Figure 5). Doerschner et al. (2011) reported a lower level of coverage for shiny surfaces than for matte surfaces because reflections appear and vanish as the object turns; the contrast on which we focus is instead between metal and glass, and the refracted image seen through a glass body is low in contrast and changes abruptly, such that fewer of its points can be followed than on a mirror-like surface, over which reflections slide coherently. Among the image statistics, the strongest cues captured differences relative to the surround: maximum object-to-surround luminance for metal and object-to-surround chroma for glass. A mirror-like surface can reflect bright light sources and the sky, thus making its brightest point brighter than the surrounding scene, whereas glass transmits the colors of the background and tends to retain their chroma. Both cues are therefore relational: they depend not on the object alone but rather on how its appearance differs from its surroundings.

Several limitations qualify the results of this research. The observer decides at some moment during the trial, and the analysis does not know when; the model reads the whole video, including the frames that are generated after the decision, which are unlikely to have contributed to it. The analysis is thus correlational, as analyses of free exploration have generally revealed that an observer who had begun to believe the object was metal may have turned it with the goal of confirming this belief. An analysis of the 2-s window before the response (Figure 7d) is the first step toward the moment of the decision. Furthermore, a more direct causal test may involve replaying one observer’s video to another observer through passive viewing and asking whether the second observer’s judgment follows that of the first observer, as active and passive viewing have been compared with regard to the perception of shape (Cornilleau-Pérès & Droulez, 1994; Wexler et al., 2001).

Material perception has largely been investigated as the interpretation of given images, but a single image can be ambiguous with respect to material. Active exploration can resolve this ambiguity by generating new views in which materials behave differently, in which context even simple image statistics can become informative with regard to the material. Our data reveal that observers who interacted with the same object generated videos that contained different types of evidence, which in turn gave rise to different judgments. Models of material perception under natural viewing therefore must explain not only how images are interpreted but also how observers generate images that resolve material ambiguity.

## Funding

T.M. was supported by a Sir Henry Wellcome Postdoctoral Fellowship from the Wellcome Trust (218657/Z/19/Z). H.T. was supported by JSPS KAKENHI (Grant Number JP25K21323) and the Naito Science & Engineering Foundation.

## Acknowledgments

For the purpose of open access, the authors have applied a CC BY public copyright license to any Author Accepted Manuscript version arising from this submission.

## Disclosures

The authors declare that they have no conflicts of interest.

## Appendix: Image statistics and models

Figure A1 illustrates how each image statistic mentioned in section 2.4 is computed on one frame and on the time series of one trial, and Table A1 presents all of these statistics, alongside their definitions. The sRGB values were linearized, and luminance and CIELAB chroma are relative to the display white. Object statistics are computed over the object’s pixels, excluding a 2-pixel-wide band inside the silhouette to ensure that the edge and frame difference energies measure the surface rather than the outline. The relational statistics are the mean, standard deviation and maximum of the surround’s luminance, its RMS contrast and its mean chroma, as well as the log ratios of object to surround values for mean luminance, RMS contrast, maximum luminance and chroma. Table A2 presents the models and reference predictors referenced in the paper. All the models are fitted (as described in section 2.5) to the other 18 observers for the AUC of an observer’s responses and without the object (to all observers’ videos of the other 39 objects) for the AUC of the disagreement test; unless stated, “statistics” indicates the 116 whole-trial statistics (object, relational and motion) of the observer’s own video.

**Figure A1.**
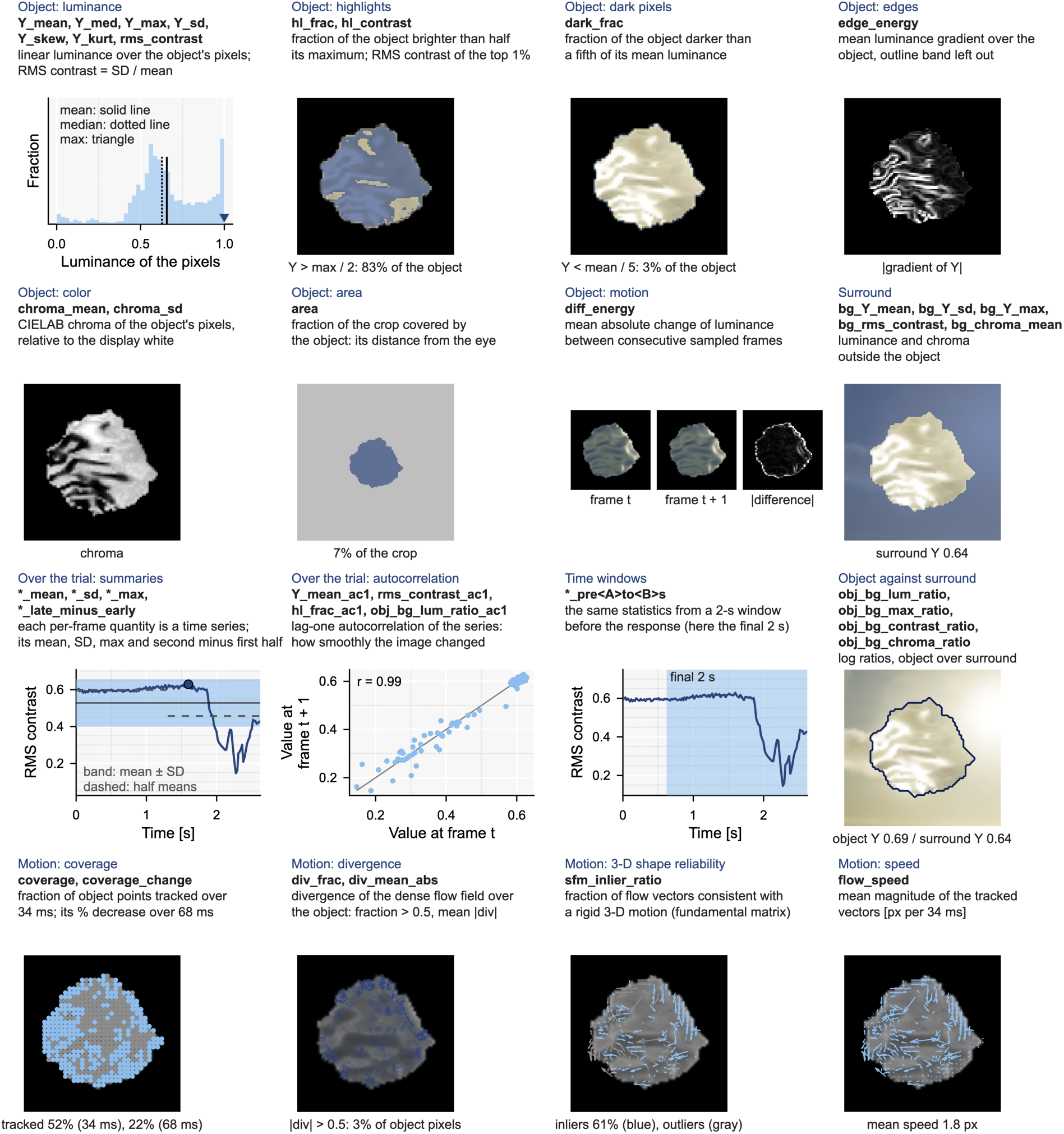
How the image statistics are computed, as illustrated on one frame of one trial (observer 5, sky, bumpy, physical metallicity 0.5, head and hand; the frame at 2.2 s of Figure 2a) and on the time series of that trial. Object statistics, on the object crop: the distribution of linear luminance over the object’s pixels, which is summarized by the mean, median, maximum, standard deviation, skewness, kurtosis and RMS contrast; the pixels brighter than half the maximum (highlight coverage) and the RMS contrast of the brightest 1%; the fraction of pixels below a fifth of the mean; the mean gradient magnitude; the mean and standard deviation of CIELAB chroma; the fraction of the crop that the object covers; and the mean absolute change in luminance between consecutive sampled frames. Relational statistics on the patch: the luminance and chroma of the surround and the log ratios of mean luminance, RMS contrast, maximum luminance and chroma between the object (outlined) and its surround. Over the trial, each per-frame quantity forms a time series, which is summarized by its mean, standard deviation, and maximum, as well as the difference between the second half and the first half; for four of these quantities, the lag-one autocorrelation is used; the time windows restrict the frames to a 2-s window before the response (the shaded span marks the final 2 s). Motion (bottom row), on one pair of frames 34 ms apart from the middle of the same trial (i.e., the motion thumbnails of the object row use the same pair): the grid points tracked between the frames (coverage, and its change when the distance is doubled), the divergence of the dense flow field, the vectors consistent with a rigid 3-D motion (inliers of a fundamental matrix) and the mean speed. Together, this set includes 15 object and 9 relational quantities × 4 summaries + 4 autocorrelations = 100 image statistics per trial, plus 4 motion quantities × 4 summaries = 16 motion statistics, accounting for a total of 116.

**Table A1.** The 116 statistics. Each of the 24 per-frame quantities is computed on every sampled frame of a video and summarized over the trial on the basis of four summaries; four quantities add the autocorrelation of their time series, 24 × 4 + 4 = 100 image statistics; the 4 motion quantities, which are computed on pairs of frames, add 4 × 4 = 16, 116 in total. The names presented in parentheses are those used in the figures.

| Statistic (the names are included in the data files) | Definition | What it captures |
|---|---|---|
| **Summaries over the trial (4 per quantity, in addition to 4 autocorrelations)** |  |  |
| <quantity>_mean, _sd, _max, _late_minus_early | Mean, standard deviation and maximum of the per-frame values over the sampled frames of the trial and the mean over the second half of the frames minus the mean over the first half. | Where the quantity sat during the trial, how much it varied, its extreme, and whether it rose or fell as the observer explored. |
| <quantity>_ac1 (for Y_mean, rms_contrast, hl_frac and obj_bg_lum_ratio) | Lag-one autocorrelation of the per-frame series. | Whether the quantity changed smoothly from one frame to the next or jumped about. |
| **Object quantities (15; over the object's pixels of the object crop, excluding a 2-pixel band inside the silhouette; Y is linear luminance relative to the white in the display)** |  |  |
| area (Object area) | Fraction of the object crop covered by the object. | How large the object appears; the crop has a fixed size, so this statistic records how close the observer brought the object. |
| Y_mean (Mean luminance) | Mean of Y over the object. | How bright the object is. |
| Y_sd (Luminance *SD*) | Standard deviation of Y over the object. | How much the brightness varies across the object. |
| Y_med (Median luminance) | Median of Y. | The typical brightness, unaffected by a few highlights. |
| Y_max (Maximum luminance) | Maximum of Y. | How bright the brightest point is. |
| Y_skew (Luminance skewness) | Skewness of the distribution of Y. | Whether a small number of very bright pixels sit on a darker body (positive) or the reverse. |
| Y_kurt (Luminance kurtosis) | Kurtosis of the distribution of Y. | How concentrated the extreme values are: sparse, intense highlights indicate a high value. |
| rms_contrast (RMS contrast) | Standard deviation of Y divided by its mean. | The contrast of the object's shading and reflections. |
| hl_frac (Highlight coverage) | Fraction of pixels brighter than half the maximum. | How much of the object is filled with bright light. |
| hl_contrast (Highlight contrast) | Standard deviation over mean of Y within the brightest 1% of pixels. | Whether the highlights are uniform or structured. |
| edge_energy (Edge energy) | Mean magnitude of the luminance gradient. | How much fine, sharp structure the object exhibits, as in mirror reflections. |
| dark_frac (Dark fraction) | Fraction of pixels below a fifth of the mean. | How much of the object is dark. |
| chroma_mean (Chroma mean) | Mean CIELAB chroma over the object. | How colorful the object is. |
| chroma_sd (Chroma *SD*) | Standard deviation of chroma. | How much the colorfulness varies across the object. |
| diff_energy (Frame difference) | Mean absolute change of Y between consecutive sampled frames over the pixels inside the object in both cases. | How much the object's image changes from one moment to the next as the observer moves. |
| **Relational quantities (9; on the patch, the surround is the patch minus the object)** |  |  |
| bg_Y_mean (Surround luminance) | Mean of Y over the surround. | How bright the background around the object is. |
| bg_Y_sd (Surround luminance *SD*) | Standard deviation of Y over the surround. | How variable the background is. |
| bg_Y_max (Surround maximum luminance) | Maximum of Y over the surround. | The brightest point of the background. |
| bg_rms_contrast (Surround RMS contrast) | Standard deviation over mean of Y in the surround. | The contrast of the background. |
| bg_chroma_mean (Surround chroma) | Mean chroma over the surround. | How colorful the background is. |
| obj_bg_lum_ratio (Object/surround luminance) | log(mean Y of the object/mean Y of the surround). | Whether the object is brighter or darker than the content that surrounds it. |
| obj_bg_contrast_ratio (Object/surround contrast) | log(RMS contrast of the object/RMS contrast of the surround). | Whether the object shows more contrast than its surround. |
| obj_bg_max_ratio (Object/surround maximum) | log(maximum Y of the object/maximum Y of the surround). | Whether the object's brightest point outshines the background. |
| obj_bg_chroma_ratio (Object/surround chroma) | log((mean chroma of the object + 1)/(mean chroma of the surround + 1)). | How colorful the object is in comparison with its surround: a glass body takes the color of what lies behind it, whereas a mirror-like body mixes the colors of the whole environment. |
| **Motion quantities (4; on pairs of frames two captures approximately 34 ms apart from a grid of points on the object every 6 pixels; the divergence fraction and the 3-D shape reliability of Doerschner et al. (2011), which are undefined in 1 to 2% of the videos, are left out)** |  |  |
| coverage (Flow coverage) | Fraction of the grid points that are tracked to the next frame (pyramidal Lucas–Kanade, kept if tracking back returns within 1 pixel). | How much of the object's image moves sufficiently coherently to be followed. |
| coverage_change (Flow coverage change) | Percent decrease of the coverage when the frame distance is doubled. | How quickly the trackable structure is lost as the object turns. |
| div_mean_abs (Flow divergence (mean)) | Mean absolute divergence of the dense flow field over the object. | Whether the image motion spreads apart or converges, as reflections do when they slide over a curved surface. |
| flow_speed (Flow speed) | Mean magnitude of the tracked vectors. | How quickly the object's image moves between frames. |

**Table A2.** Models and reference predictors.

| Model | Input for each trial | Fitting | Test | Section |
|---|---|---|---|---|
| Physical metallicity | The physical metallicity of the trial (0 to 1). | Nothing (a reference predictor). | AUC for each observer's responses. | 3.1, 3.2 |
| Other observers | Proportion of the other 18 observers who responded "metal" on the same trial (lower bound of the ceiling) or of all 19 including the observer (upper bound). | Nothing (a reference predictor). | AUC; the ceiling for a model that uses nothing that is specific to the observer. | 3.1, 3.2 |
| Model | Input for each trial | Fitting | Test | Section |
| Own-video statistics | The 63 object, 37 relational and 16 motion statistics of the observer's own video, whole trial. | As in section 2.5 (C = 0.1). | Leave-one-observer-out AUC, by condition; AUC of the disagreement test against the permutation null. | 3.2, 3.4 |
| Statistics of the single image | The 23 per-frame quantities (14 object, 9 relational) of the image that the observer saw in the condition without exploration. | As in section 2.5, within that condition (C = 0.1); also the per-frame means of the moving videos, which are fitted to the other 18 observers' moving videos and applied to the image. | Leave-one-observer-out AUC. | 3.2 |
| 2-s window before the response | The same statistics from a 2-s window of the video that ends 0, 1, 2, 3, 4, 5 or 6 s before the response; only the videos that cover the window are used. The final window was also computed by reference to videos that were 3 s long or longer only, including the videos of the window prior to it. | As in section 2.5 (C = 0.1). | AUC of the disagreement test against the permutation null. | 3.4 |
| Duration only | The logarithm of the number of frames. | As in section 2.5 (C = 0.1). | AUC of the disagreement test. | 3.4 |
| Single statistics | One statistic alone, which is standardized within the observer. | Nothing. | AUC for each observer's responses. | 3.3 |
| Image statistics without motion | The 100 object and relational statistics, without the 16 motion statistics. | As in section 2.5 (C = 0.1). | Leave-one-observer-out AUC; AUC of the disagreement test. | 3.2 |
| Pairs by the difference between the videos | The own-video statistics, whole trial; the disagreeing pairs are split into quarters by the Euclidean distance between the two videos' scores on the 40 principal components of the standardized statistics (95% of their variance). | As in section 2.5 (C = 0.1). | AUC of the disagreement test of each quarter by reference to its permutation null (quarter boundaries fixed). | 3.4 |
| Permutation null | The own-video model's predicted metallicity with the responses shuffled among observers within each trial (300 shuffles). | Nothing. | Distribution of the AUC of the disagreement test under the null. | 3.4 |

## Notes

### Competing Interest Statement

The authors have declared no competing interest.

